# Lotka-Volterra Equations in the Presence of Mimicry

**DOI:** 10.1101/2021.03.25.436931

**Authors:** Inavamsi Enaganti, Bud Mishra

## Abstract

This paper studies a novel framework for understanding toxicity and nutritional value of a prey to a predator with utility. Furthermore, it introduces a novel notion of prey mimicry in a multi predator - multi prey system. In addition, the paper also analyzes the dynamics and the stability of equilibrium that arise as a result of the one or more predators’ inability to distinguish between certain prey. The paper is motivated by the mimicry in the context of adversarial chase involved in the evolution of multiple viral variants in the presence of multiple vaccines – possibly, in a population stratified by a diverse HLA background.

## 1. INTRODUCTION

The dynamics between predator and prey is a fascinating subject. This subject has been extensively studied and formalized. We introduce the notion of prey toxicity and mimicry and study the resulting dynamics. This can easily be extended to other fields and sciences. A classic example would be the dynamics behind counterfeit currency which is essentially a toxic mimic to real currency. Other examples are that of the CAPTCHA (Turing test) vs. fake bots or vaccines against a mutating virus which are becoming more and more relevant today. One of these examples has been elaborated in Appendix C.

We assume a simple system with multiple predators and prey. Each prey species can either be toxic or nutritious to a predator. A rational predator will always consume the nutritious prey and avoid the toxic prey. The predator needs to be able to distinguish between the different prey species.

To distinguish between prey of different species, the predator uses the signal sent by the prey. The signal can be anything from color, smell, sound or even a dance pattern. So a red toxic species can easily be distinguished from a blue nutritious species. But in the case where both prey send the same signal to a predator, they cannot be distinguished by the predator. The predator essentially treats them as a single prey and this can be considered as mimicry. [1]

Interestingly, mimicry depends on the point of view. For example, two prey can both be red in color but produce different sounds. So a predator that hunts using just visual signals will not be able to distinguish between the two. Whereas the predator that hunts using acoustic signals will be able to distinguish between the two prey. Furthermore, a prey can be toxic to certain predators but nutritious to others. The toxicity and nutritional values is encoded as utility of the prey to the predator. We specifically explore the dynamics of this example in further sections.

The concept of mimicry becomes trivial without the presence of toxicity. To sum up, we aim to study the interesting dynamics of a predator prey system with the presence of mimicry and toxicity. This has been shown by generalizing the standard Lotka-Volterra equations. [2]

## 2. GENERALIZED LOTKA-VOLTERRA EQUATIONS FOR MULTI PREDATOR - MULTI PREY MODEL

In general for a multi predator-multi prey model we consider the following assumptions.

1. The prey population finds ample food at all times.
2. The food supply of the predator population depends entirely on the size of the prey population.
3. The rate of change of population is proportional to its size.
4. During the process, the environment does not change in favour of one species, and genetic adaptation is inconsequential.
5. Predators have limitless appetite.
6. In the absence of prey the predators exponentially decay.
7. There is no competition between the prey.
8. Every prey species has a fixed utility for every predator species.

We consider n prey where each prey is denoted by *x*_*i*_, *i ∈* {1…*n*}. We consider m predator where each predator is denoted by *y*_*j*_, *j ∈* {1…*m*}.

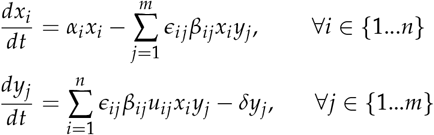

Where,

- ϵ _*i j*_*∈* {0, 1} is an indicator variable indicating whether *x*_*i*_ is a prey for predator *y*_*j*_.
- α _*i*_ *>* 0 corresponds to growth rate of prey *x*_*i*_
- *β* _*ij*_ *>* 0 denotes the number of interactions or rate of predation between prey *x*_*i*_ and predator *y*_*j*_
- *u*_*ij*_ denote the utility of prey *x*_*i*_ to predator *y*_*j*_. This can be scaled as required
- *δ* _*j*_ *>* 0 denote the loss/death rate of predator *y*_*j*_

If *u*_*i j*_ *>* 0 then prey *x*_*i*_ is nutritious to predator *y*_*j*_ and is beneficial. If *u*_*ij*_ *<* 0 then prey *x*_*i*_ is toxic to predator *y*_*j*_ and is fatal.

If a predator *y*_*j*_ can distinguish between a toxic *x*_*t*_ and a nutritious *x*_*n*_. Then ϵ_*n*__*j*_ *>* 0, while ϵ_*t*__*j*_ = 0. Optimally, the predator should choose to not consume the toxic prey. The interesting case is when the nutritious prey *x*_*n*_ mimics toxic prey *x*_*t*_. If a prey *x*_*i*_ is toxic to predator *x*_*j*_then we consider that ϵ_*ij*_ = 0. This gets interesting when the predator cannot distinguish between a toxic prey and a nutritious prey. This is also known as Batesian mimicry.

## 3. MIMICRY IN A SIMPLE TWO PREDATOR - TWO PREY MODEL

We consider two prey A and B. We also consider two predators P and Q. We assume the same exponential growth rate for both prey (*α* = *α*_*A*_ = *α* _*B*_), the same rate of interactions for every pair of predator and prey(*β* = *β* _*AP*_ = *β* _*AQ*_ = *β* _*BP*_ = *β* _*BQ*_) and the same exponential loss/death rate for both predators (*δ* = *δ* _*P*_ = *δ* _*Q*_)

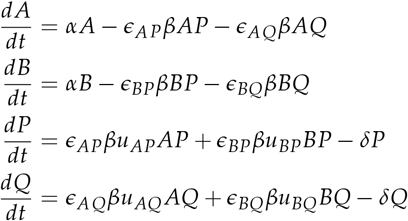

### A. Single Predator - Two Prey

Consider an ecosystem with only one type of predator P (*Q* = 0). A is nutritious to predator P (*u*_*Ap*_ *>* 0) while prey B is toxic to predator P (*u*_*BP*_ *<* 0).

**Case 1:**If A and B can be distinguished by the predator P.

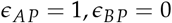

**Case 2:**If A mimics toxic B and cannot be distinguished by the predator P.

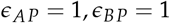

**Case 2.1:** If *u*_*BP*_ A + *u*_*BP*_B ≥ 0

This can be treated as single predator single prey model with the utility of the single prey being the expected utility which is 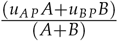. As the predator cannot distinguish between the two prey species it sees a single signal which can be attributed to an expected reward or utility.

Given that the growth rate of A and B is the same, if *β* _*AP*_ *≠ β* _*BP*_ then the prey species with higherV rate of interaction goes extinct. This is a standard result of one predator - two prey model.

**Case 2.2:** If *u*_*AP*_ A + *u*_*BP*_B < 0

The rate of change of predators is negative. Thus both predators decay and die out while the prey under no predation grow exponentially.

**Equilibrium** In an equilibrium or a fixed point of the system all the derivatives are zero. [3]

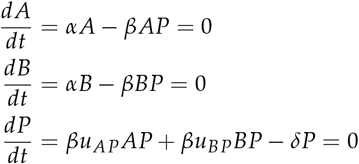

The fixed points are (*A, B, P*) = (0, 0, 0) and 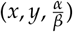 such that *u*_*AP*_*x* + *u*_*BP*_*y* = *δ/ β*. To check for stability of the fixed points we check whether the real part of all eigen values of the Jacobian are negative.

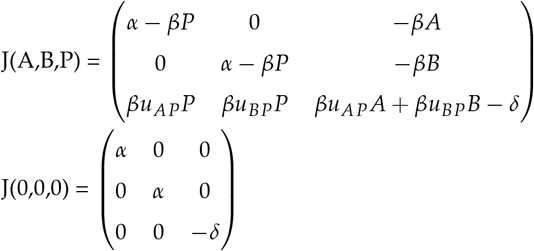

α is an eigenvalue and is positive hence this is not a stable fixed point. This shows that (0,0,0,0) is not stable even in the generalised version.

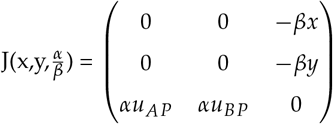

We see that *λ*^3^ = *λ*(*− α δ*). One of the eigenvalues is 0 and the other two are imaginary conjugates. This results in a center equilibrium with periodic orbits. [3]

### B. Two Predator - Two Prey

We look at a few examples of mimicry in this model.

- Prey A and B are distinctly identifiable by both the predators P and Q.
- Prey A and B can be distinguished by predator P but not by Q (or vice versa).
- Prey A and B can’t be distinguished by either predator P or Q.

When a predator *y*_*j*_ can distinctly identify a prey *x*_*i*_ and the corresponding utility *u*_*ij*_ is negative then clearly ϵ_*i j*_ = 0.

In the trivial case where both prey have positive utility to both predators. The model follows a standard two predator-two prey model. In the case that both prey have negative utility to both predators. Both predators die out.

Note: Table 1, 2, 3 show a few cases for each condition. The remaining cases can be obtained with a change in labels of shown cases. Consider all unmentioned ϵ values to be 1 by default.’ +’ implies positive utility where u > 0. Similarly’’ implies negative utility where u < 0.

**Table 1.**
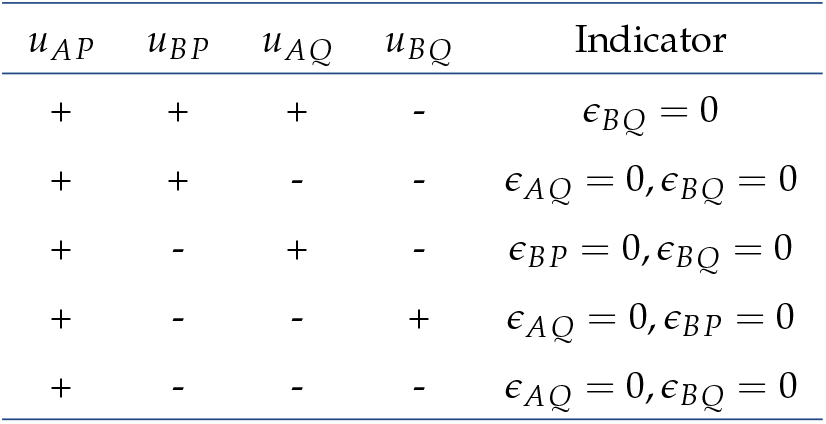
Prey A and B are distinctly identifiable by both the predators P and Q.

**Table 2.** Prey A and B can be distinguished by predator P but not by Q (or vice versa).

| $u_{AP}$ | $u_{BP}$ | $u_{AQ}$ | $u_{BQ}$ | Indicator |
| --- | --- | --- | --- | --- |
| + | + | + | - | Depends on expected utility |
| + | + | - | - | $\epsilon_{AQ} = 0, \epsilon_{BQ} = 0$ |
| + | - | + | - | Not fixed |
| + | - | - | + | Not fixed |
| + | - | - | - | $\epsilon_{BP} = 0, \epsilon_{AQ} = 0, \epsilon_{BQ} = 0$ |

**Table 3.** Prey A and B cannot be distinguished by either predator P or Q.

| $u_{AP}$ | $u_{BP}$ | $u_{AQ}$ | $u_{BQ}$ | Indicator |
| --- | --- | --- | --- | --- |
| + | + | + | - | Depends on expected utility |
| + | + | - | - | $\epsilon_{AQ} = 0, \epsilon_{BQ} = 0$ |
| + | - | + | - | Depends on expected utility |
| + | - | - | - | $\epsilon_{AQ} = 0, \epsilon_{BQ} = 0$ |

The indicator variables can be interpreted as following.

1. If ϵ _*i j*_ = 0 and ϵ_*i*_*′* _*j*_ = 0 then predator *y*_*j*_ will go extinct. The final model is essentially a one predator - one prey model.
2. If ϵ _*i j*_ = 0 and ϵ_*ij*_*′*= 0 then prey *x*_*i*_ will grow exponentially. The final model is essentially a one predator - one prey model.
3. If ϵ _*i j*_ = 0 and ϵ_*i*_*′* _*j*_*′*= 0 and *i* ≠ *i′* and *j* ≠ *j′*, then the model is essentially two independent one predator - one prey model.

4. If predator *y*_*j*_ cannot distinguish between *x*_*i*_ and *x*_*i*_*′* then

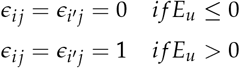

Where 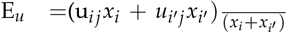 is the expected utility predator *y*_*j*_ gets from the mimicry ring.

### C. Special Cases of Two Predator - Two Prey

The interesting situation occurs in Table 2 case 3 and 4 labelled as’Not fixed’.

We take a look at case 3 defined as follows. Prey A and B can be distinguished by predator P but not by Q.

In case 3, the utilities are as follows. *u*_*AP* *>*_ 0, *u*_*BP*_ *<* 0, *u*_*AQ*_ *>* 0, *u*_*B*__*Q*_ *<* 0.

In case 4, the utilities are as follows. *u*_*AP*_ *>* 0, *u*_*BP*_ *<* 0, *u*_*AQ*_ *<* 0, *u*_*B*__*Q*_ *>* 0

Predator P can distinguish between prey A and B and thus will never choose to consume B as utility is negative. On the other hand predator Q has no choice but to consume whatever prey it encounters to survive.

We assume the following ϵ’s. It trivially reduces in other cases as seen before. ϵ_*AP*_ = 1, ϵ_*BP*_ = 0, ϵ_*AQ*_ = 1, ϵ_*B* *Q*_ = 1

In a equilibrium or a fixed point of the system all the derivatives are zero. The trivial fixed point is (*A, B, P, Q*) = (0, 0, 0, 0). The other fixed point is when *P* = 0 and thus exactly the same as Case 2 of one predator - two prey model mentioned before.

It is impossible for both predator and prey species to all be alive in an equilibrium, this is because A will be predated more than B as long as P is non zero.

## 4. GENERALIZING MIMICRY IN TWO PREDATOR - TWO PREY MODEL

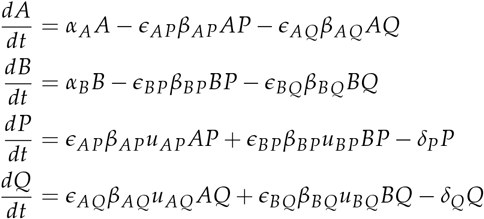

**Equilibrium** There are multiple fixed points or equilibrium but we want to focus on the equilibrium where none of the prey and none of the predators go extinct.

There is only one fixed point at which all *A, B, P, Q* ≠ 0. The fixed point is (a,b,p,q) where

- 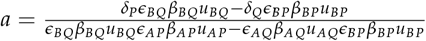
- 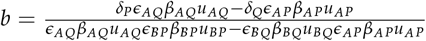
- 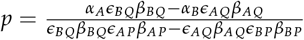
- 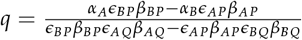

The Jacobian will be as follows. *J*(*A, B, P, Q*)

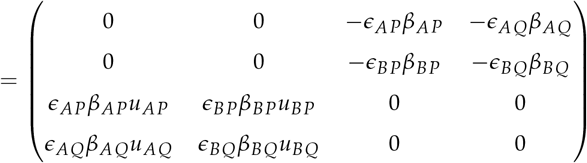

We consider the special cases we saw in the previous section. All other cases devolve to simpler models.

Prey A and B can be distinguished by predator P but not by Q. The utilities are as follows. We consider static indicator variables. Thus ϵ_*AP*_ = 1, ϵ _*BP*_ = 0, ϵ_*AQ*_ = 1, ϵ_*B*__*Q*_ = 1

The fixed point becomes (a,b,p,q) where

- 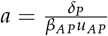
- 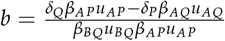
- 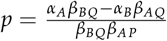
- 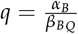

**Special Case 1** *u*_*AP* *>*_ 0, *u*_*BP*_ *<* 0, *u*_*AQ*_ *>* 0, *u*_*B*__*Q*_ *<* 0 Conditions for equilibrium

- *α* _*A*_ *β* _*BQ*_ > *α* _*B*_ *β* _*AQ*_
- *δ* _*P*_ *β* _*AQ*_ *u* _*AQ*_ > *δ* _*Q*_ *β* _*AP*_ *u* _*AP*_

**Special Case 2** *u*_*AP*_ *>* 0, *u*_*BP*_ *<* 0, *u*_*AQ*_ *<* 0, *u*_*B*__*Q*_ *>* 0 Conditions for equilibrium

- *α* _*A*_ *β* _*BQ*_ > *α* _*B*_ *β* _*AQ*_

For both special cases the Jacobian is as follows.

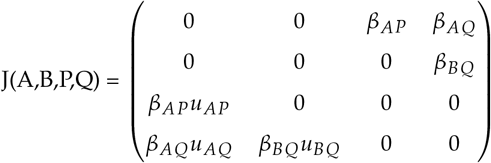

There exists a positive eigenvalue for the above matrix if *u*_*AP*_*u*_*AQ*_*u*_*BQ*_ *<* 0. See supplemental material for proof. Thus this is an unstable equilibrium. [3]

We can conclude that there is no stable equilibrium in a two predator - two prey model where each predator has one toxic and one non-toxic prey when one predator can distinguish between the two while the other cannot.

**Example:** We consider an equilibrium/fixed point. By perturbing the parameters we see that it is not a stable equilibrium.

This has been shown in Fig.4 and Fig5.

**Fig. 1.**
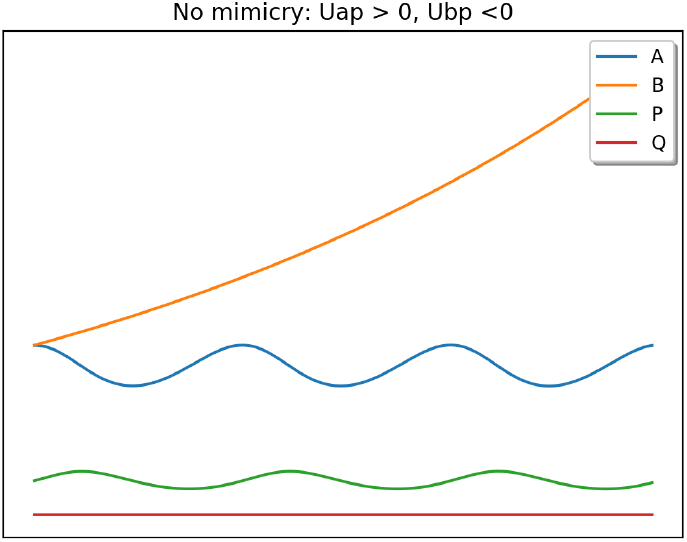
Prey B grows exponentially while A and P follow the standard single predator - single prey model.

**Fig. 2.**
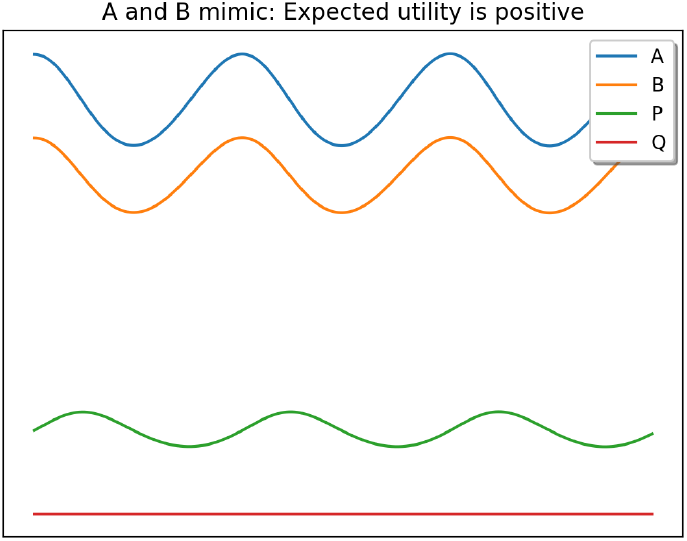
Expected utility is positive. Effectively a one predator - one prey model.

**Fig. 3.**
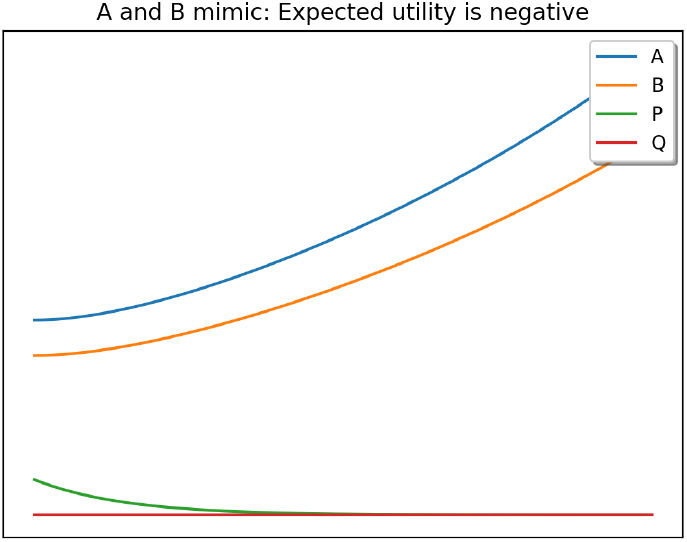
Expected utility is negative. Predator dies out and then prey grows exponentially.

**Fig. 4.**
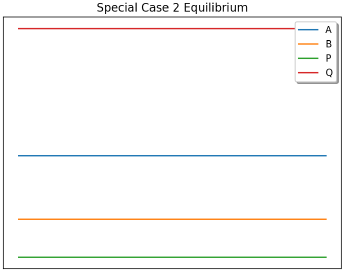
Special Case 2: Equilibrium Fixed point according to α_*A*_β_*B*__*Q*_ *>* α _*B*_ β_*BQ*_.

**Fig. 5.**
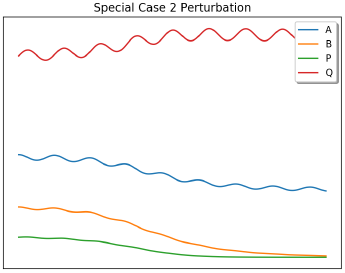
Special Case 2: Unstable equilibrium with small perturbation α_*A*_β _*BQ*_*>* α _*B*_β_*BQ*_.

**Fig. 6.**
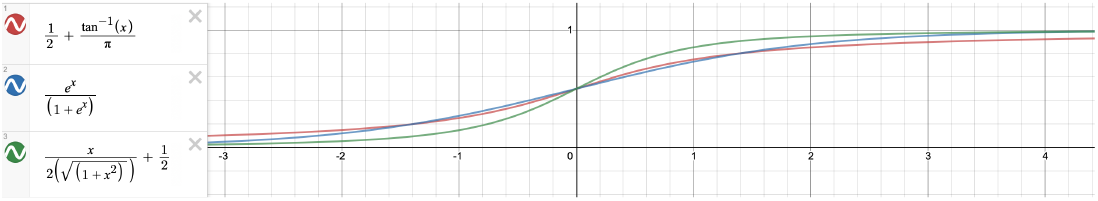

**Fig. 7.**
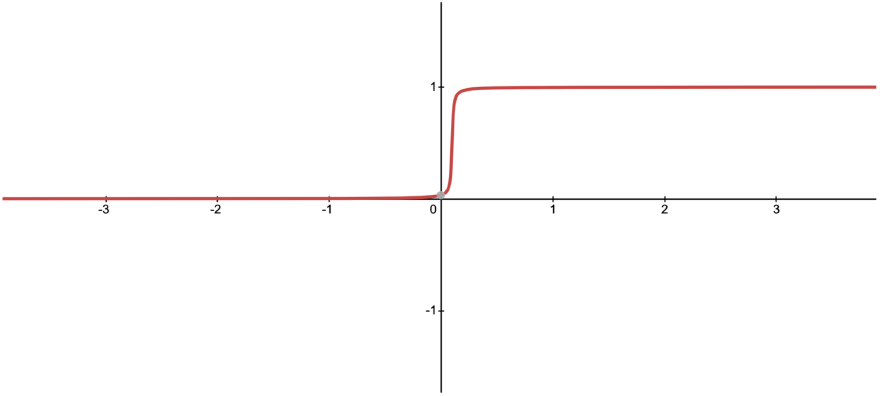
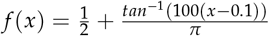

## 5. CONCLUSION

We have presented a formal framework for analysing dynamics of the signaling game between predator and prey using utility.

We have also shown that there is no stable equilibrium in a two predator-two prey system in the presence of mimicry of two prey in front of a predator.

## 6. DISCUSSION

We have assumed a non evolving system where the predator does not continually learn and evolve as well, although at a slower pace. A possible solution to the learning capability of a predator is a dynamic indicator variable which has been discussed below.

This will be discussed in a following paper which treats signals and utility as related quantities and allows for their evolution through the natural selection of the predator.

### A. Generalizing to m predator n prey mimicry

When a predator *y*_*j*_ can distinctly identify a prey *x*_*i*_ and the corresponding utility *u*_*ij*_ is negative then clearly β_*i j*_ = 0. On the other hand if a predator *y*_*j*_ cannot distinguish between some prey species *x* _*s*_, *s ∈ S ⊆* {1…*n*}. The predator views this chunk as a single unit and the utility will be the expected utility from the entire chunk. The expected utility is 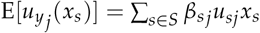.

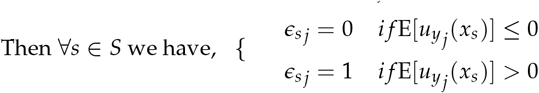

The problem starts here as the expected utility is dynamically changing with time as other predators interact with specific subsets of the population.

#### B. Dynamic Indicator Variable

ε ϵ [0,1]

We consider a dynamic epsilon value that continuously changes with the expected utility. This will effectively allow the predator to be rational and choose a signal only if it is getting a positive utility from it. And similar to natural predators as utility approaches 0, the indicator variable approaches 0. For this we use an activation function

We look at some standard activation functions.

But these functions still have significant indicators when utility is negative. Thus we can scale and translate these functions to behave like the following.

As utility becomes negative the predator will drastically reduce the amount of prey consumed. We consider the small value to be the exploration parameter to constantly check the utility it would receive.

Thus for Mimicry our indicator variable will look like the following if A and B are not distinguishable by Q.

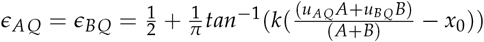

##### C. Practical Example

Two competing companies A and B have people visiting their websites. Each company invests money into advertising/services to those that visit their website. Both companies create bots to bring down their competitors. It is beneficial is the recipient was human, but resources wasted if the recipient was a bot.

A bot which is the malicious agent will try to solve the Turing test to pass off as a human. The websites will get bad utility from a bot and thus create better Turing test which are harder for bots to crack. This resembles a co-evolutionary chase between predator and prey. The bot is trying to mimic the human introducing the element of mimicry. This perfectly mirrors two predators(Companies) consuming prey(visitors). Some prey are toxic(bots) and some nutritious(humans). Their ability to distinguish between the signals(Captcha/Turing test) changes with time as they learn. If we know the dynamics one can easily analyze the amount of money a company must spend on creating bots for competitors or how much one must invest into a Turing Test.

## 7. SUPPLEMENTAL MATERIAL

**Proof of claim** : There exists a positive eigenvalue for the below matrix if *u*_*AP*_*u*_*AQ*_*u*_*BQ*_ *<* 0.

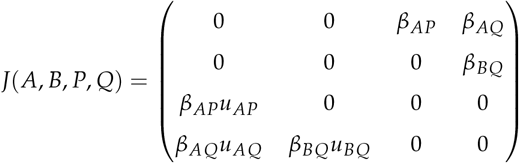

The characteristic equation is

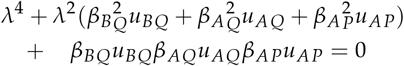

This is of the form *ax*^2^ + *bx* + *c* = 0 where

- *x* = *λ* ^2^
- *a* = 1
- 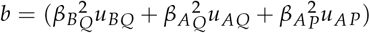
- *c* = *β* _*BQ*_ *u*_*BQ*_ *β* _*AQ*_ *u*_*AQ*_ *β* _*AP*_ *u*_*AP*_

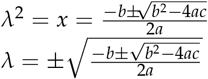

We consider the following root. 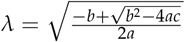

Since *u*_*AP*_*u*_*AQ*_*u*_*BQ*_ *<* 0 it follows that 4*ac <* 0. Thus 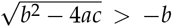. Which results in 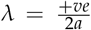 which is a positive number.

Thus we have found the required positive eigenvalue(*λ*).

